# Malaria mosquitoes acquire and allocate cattle urine to enhance life history traits

**DOI:** 10.1101/2020.08.24.264309

**Authors:** Mengistu Dawit, Sharon R. Hill, Göran Birgersson, Habte Tekie, Rickard Ignell

## Abstract

Nutrient acquisition and allocation integrate foraging and life-history traits in insects. To compensate for the lack of a particular nutrient at different life stages, insects may acquire these through supplementary feeding on *e.g.*, vertebrate secretions, in a process known as puddling. The mosquito *Anopheles arabiensis* emerges undernourished, and as such, requires nutrients for both metabolism and reproduction. Host-seeking and blood-fed *An. arabiensis* are attracted to the natural and synthetic odour of cattle urine, which signals a source of nutrients, but not the presence of a host or oviposition site. Females actively imbibe cattle urine, and its main nitrogenous compound, urea, and allocate these resources according to life history trade-offs to flight, survival or reproduction, as a function of physiological state. As a consequence, this behaviour affects vectorial capacity by increasing daily survival and vector density, and thus should be considered in future models. Future vector management strategies are discussed.

## Introduction

Acquisition and allocation of nutrients integrate foraging and life-history traits in insects (Boggs, 2009; Molleman, 2010; Raubenheimer et al., 2009). Insects are capable of selecting and acquiring diets, and of compensatory feeding, in response to food availability and need for nutrients (Boggs, 2009; Raubenheimer et al., 2009). Allocation of nutrients is dependent on life-history processes, and may result in different needs of diet quality and quantity at different life stages of the insect (Boggs, 2009; Molleman, 2010). To compensate for the lack of a particular nutrient, insects may acquire these through supplementary feeding on *e.g.*, mud, various excrements and secretions of vertebrates, and carrion, in a process referred to as puddling (Molleman, 2010). Although mainly described for various butterfly and moth species, puddling also occurs in other insect orders, where attraction to and feeding on these types of resources has a significant effect on fitness and other life-history traits (Bänziger et al., 2009; Hendrichs et al., 1993; Molleman, 2010; Plotkin and Goddard, 2013; Shen et al., 2009). The malaria mosquito, *Anopheles gambiae sensu lato*, ecloses as an ‘undernourished’ adult (Van Handel, 1965), and, as such, puddling may play an important role for its life-history traits, but is a behaviour that so far has been overlooked. The inclusion of puddling as a means to enhance nutrient intake in this important vector requires attention, as this may have important epidemiological consequences.

Due to low caloric reserves carried over from the larval stage and a low efficiency of blood meal utilization (Briegel and Horler, 1993), adult female malaria mosquitoes are limited in their nitrogen intake. Female *An. gambiae s.l.* often compensate for this by taking multiple blood meals (Klowden and Briegel, 1994; Norris et al., 2010), thereby putting more people at risk of contracting disease. Alternatively, mosquitoes could use supplementary feeding on vertebrate excretions to obtain nitrogenous compounds to enhance fitness and flight mobility, as demonstrated for other insects (Molleman, 2010). In this regard, the strong and differential attraction of one of the sibling species within the *An. gambiae s.l.* species complex, *An. arabiensis*, to fresh and aging cattle urine (Kweka et al., 2009; 2011; Mahande et al., 2010), is intriguing. Cattle urine is a resource rich in nitrogenous compounds, with urea making up 50-95 % of the total nitrogen in fresh urine (Dijkstra et al., 2013; Kilande et al., 2016). As cattle urine ages microbes make use of these resources, thereby reducing the complexity of nitrogen-containing compounds within 24 h (Kilande et al., 2016). With the rapid increase in ammonia, correlating with the decline in organic nitrogen, alkalophilic microbes, many of which produce compounds toxic to mosquitoes, thrive (Kilande et al., 2016), which may be a lead cause of why female *An. arabiensis* are preferentially attracted to urine aged for 24 h or less (Kweka et al., 2011; Mahande et al., 2010).

In this study, we assessed whether host-seeking and blood-fed *An. arabiensis* can acquire nitrogenous compounds, including urea, through urine puddling. Next, we conducted a series of experiments to assess how female mosquitoes allocate this potential nutrient resource to enhance survival, reproduction and further foraging. Finally, we assessed whether the odour of fresh and aging cattle urine provides a reliable cue for host-seeking and blood-fed *An. arabiensis* in their search for this potential nutrient resource, and identified the chemical correlates underlying the observed differential attraction. The synthetic odour blend of volatile organic compounds (VOCs) identified in 24 h aged urine was further evaluated under field conditions, expanding on the results obtained under laboratory conditions, and demonstrating the efficacy of cattle urine odour to attract mosquitoes of different physiological states. The obtained results are discussed in the context of potential epidemiological consequences.

## Results

### Host-seeking and blood-fed mosquitoes feed on urine and urea

To assess whether *An. arabiensis* are able to acquire urine, and its main source of nitrogen, urea, through direct feeding, 4-day post-emergence (dpe) host-seeking and blood-fed females were given access to these diets over a period of 48 h in a feeding assay (Fig. 1A). Both host-seeking and blood-fed females imbibed significantly larger volumes of sucrose than any of the other diets or water (F_(5,426)_ = 20.15, p < 0.0001 and F_(5,299)_ = 56.00, p < 0.0001, respectively; Fig. 1BC). In addition, host-seeking females fed less on 72 h aged urine compared with 168 h aged urine (Fig. 1B). When provided with diets containing urea, host-seeking females imbibed a significantly larger volume of 2.69 mM urea compared with all other concentrations and water, while not differing from 10 % sucrose (F_(10,813)_ =15.72, p < 0.0001; Fig. 1D). This differed from the response of blood-fed females, which generally imbibed significantly larger volumes of urea-containing diets compared to water, although imbibing significantly smaller volumes compared to 10 % sucrose (F_(10,557)_ = 78.35, p < 0.0001; Fig. 1E). Moreover, when comparing between the two physiological states, blood-fed females imbibed more urea at the lowest concentration than their host-seeking counterparts, while these females imbibed similar amounts at higher concentrations (F_(1,953)_ = 78.82, p < 0.0001; Fig. 1FG). While the volume of intake of the urea-containing diets appeared to have an optimum (Fig. 1DE), females of both physiological states were able to regulate the amount of urea imbibed over the full range of urea concentrations in a log-linear fashion (Fig. 1FG). Similarly, mosquitoes appear to control their intake of nitrogen by regulating the volume of urine imbibed, as the amount of nitrogen in the urine is reflected in the volume imbibed (Fig. 1BC and Fig. 1B inset).

**Figure 1.**
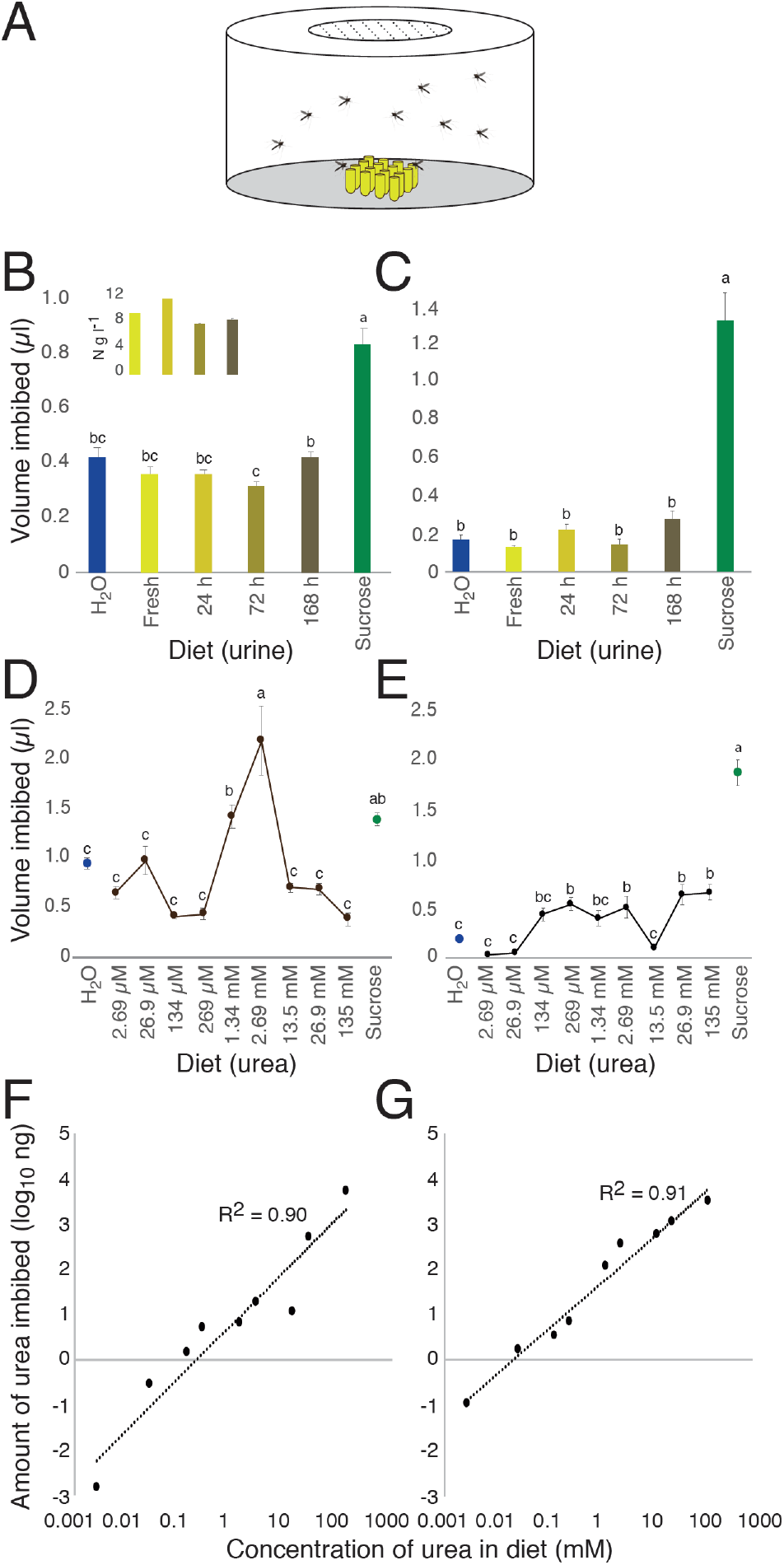
Cattle urine and urea imbibed by host-seeking and blood-fed female *Anopheles arabiensis*. Female mosquitoes were provided with diets consisting of fresh and aged cattle urine, various concentrations of urea, sucrose (10 %) and distilled water (H_2_O) in a feeding assay (A). Host-seeking (B) and blood-fed (C) females imbibed larger volumes of sucrose than any of the other diets tested. Note that host-seeking females imbibed less 72 h aged cattle urine than 168 h aged cattle urine (B). The average total nitrogen content of the urine (± standard deviation) is represented in the inset. Urea was imbibed by host-seeking (D, F) and blood-fed (E, G) females in a dose-dependent manner. The mean volume imbibed (D, E) with different letter designations are significantly different from one another (one-way analysis of variance with a Tukey’s *post hoc* analysis; p < 0.05). Error bars represent the standard error of the mean (B-E). The straight dotted lines represent the log-linear regression lines (F, G).

### Urine and urea affect the survival of malaria mosquitoes

To assess the role of urine and urea on the survival of host-seeking and blood-fed mosquitoes, females were fed on all four ages of urine and a range of urea concentrations, as well as on distilled water and 10 % sucrose as controls (Fig. 2A). This survival analysis revealed that diet had a significant impact on the overall survival rate of host-seeking females (urine: χ^2^ = 108.5, df = 5, p < 0.0001; urea: χ^2^ = 122.8, df = 5, p < 0.0001; Fig. 2BC) and blood-fed females (urine: χ^2^ = 93.0, df = 5, p < 0.0001; urea: χ^2^ = 137.9, df = 5, p < 0.0001; Fig. 2DE). In all experiments, females feeding on urine, urea and water had a significantly reduced survival compared to those provided with sucrose as a diet (Fig. 2B-E). Host-seeking females feeding on fresh and aged urine exhibited differential survival, with those feeding on 72 h aged urine (p = 0.016) having the lowest probability of survival (Fig. 2B). Moreover, host-seeking females fed on 135 mM urea survived longer than on the water control (p < 0.04) (Fig. 2C). Blood-fed females survived longer when fed on fresh and 24 h aged urine compared with water (p = 0.001 and p = 0.012, respectively; Fig. 2D), while those fed on 72 h aged urine survived for a shorter time than those fed on fresh and 24 h aged urine (p < 0.0001 and p = 0.013, respectively; Fig. 2D). When fed on 135 mM urea, blood-fed females survived longer than all other concentrations of urea and water (p < 0.013; Fig. 2E).

**Figure 2.**
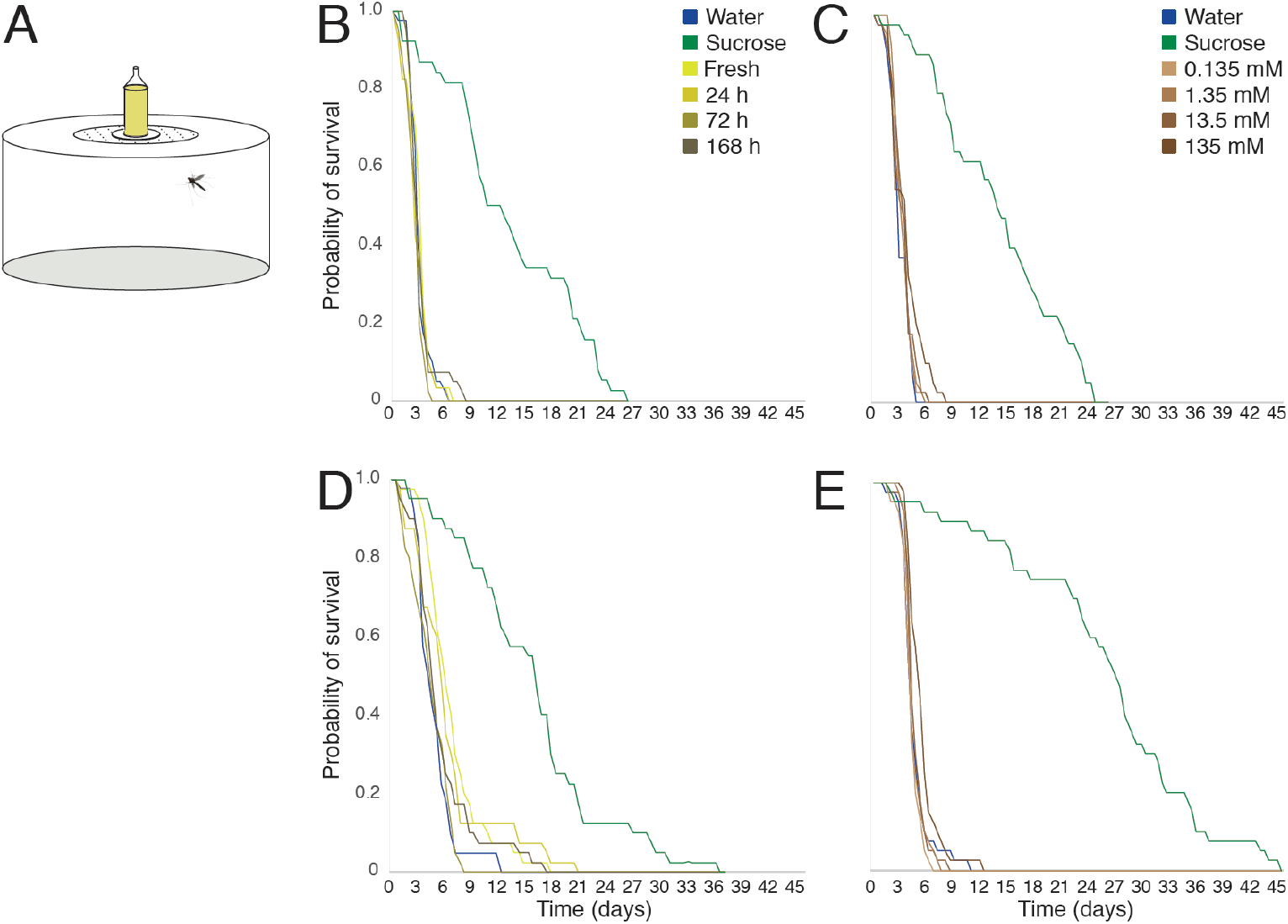
Survival of host-seeking and blood-fed female *Anopheles arabiensis* fed on cattle urine and urea. Female mosquitoes were provided with diets consisting of fresh and aged cattle urine, various concentrations of urea, sucrose (10 %) and distilled water (H_2_O) in a bio-assay (A). The survival of individual host-seeking (B, C) and blood-fed (D, E) mosquitoes was recorded every 12 h, until all females fed on urine (B, D) and urea (C, E), as well as the controls, sucrose and water, had died.

### Flight behaviour is affected by urine and urea diet

The overall distance and number of bouts, as determined in a flight mill assay over a 24 h period, differed between host-seeking and blood-fed mosquitoes, with blood-fed mosquitoes displaying less flight activity overall (Fig. 3). Host-seeking mosquitoes provided with fresh and aged urine, or sucrose and water, displayed varying flight patterns (Fig. 3), with females fed on fresh urine being more active at dawn, while those fed on 24 h and 168 h aged urine displaying predominantly daytime activity. Female mosquitoes provided with either sucrose or 72 h aged urine demonstrated activity throughout the 24 h period, whereas those provided with water were more active during mid-scotophase. Mosquitoes fed on sucrose demonstrated the highest levels of activity during the late night and early morning, while those that imbibed 72 h aged urine decreased activity steadily over the 24 h (Fig. 3).

**Figure 3.**
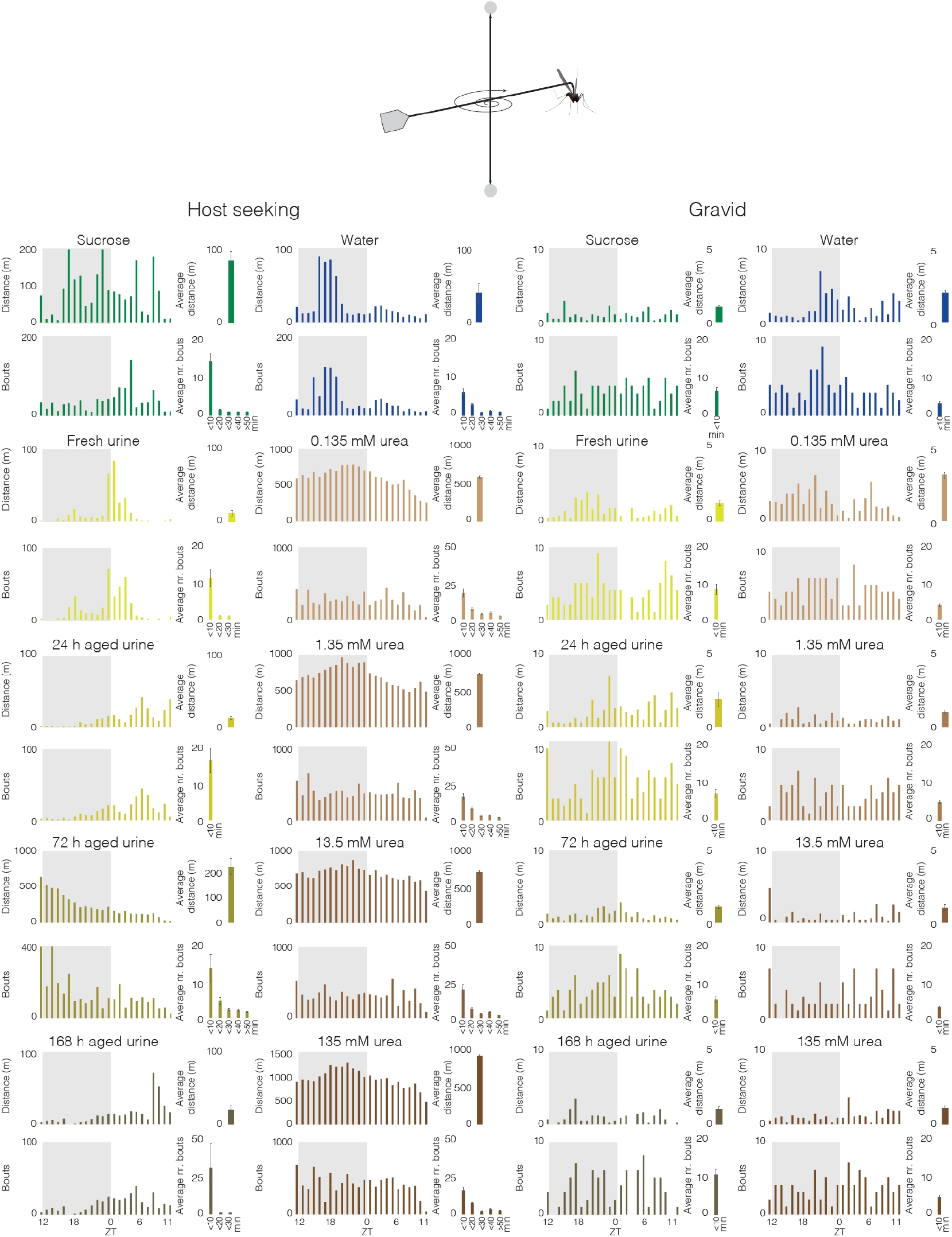
Flight performance of host-seeking and blood-fed female *Anopheles arabiensis* fed on cattle urine and urea. Female mosquitoes fed on diets consisting of fresh and aged cattle urine, various concentrations of urea, sucrose (10 %) and distilled water (H_2_O) were tethered to a horizontal, free-spinning arm in a flight mill assay (top). The overall distance and number of bouts flown per hour over 24 h (scotophase: grey; photophase: white) were recorded for each diet for both host-seeking (left) and blood-fed (right) females. The average distance and average number of bouts are shown to the right of the diurnal activity plots. Error bars represent the standard error of the mean. See the main text for statistical analysis.

In general, the overall bouts of flight activity by host-seeking females followed a similar pattern to that of the distance flown over the 24 h period. The diet imbibed significantly affected the average distance flown (F_(5, 138)_ = 28.27, p < 0.0001), with host-seeking females having imbibed 72 h aged urine flying significantly longer distances than all other diets (p < 0.0001), and sucrose-fed mosquitoes flying longer distances than those fed on fresh (p = 0.022) and 24 h aged urine (p = 0.022). In contrast to the flight activity patterns described for the urine diets, host-seeking females fed on urea demonstrated continuous flight activity over the course of the 24 h period with a peak of activity during the second half of scotophase (Fig. 3). While the pattern of activity was similar, host-seeking females fed on urea significantly increased the average distance flown depending on the concentration imbibed (F_(5, 138)_ = 1310.91, p < 0.0001). Host-seeking females feeding on any concentration of urea tested flew longer distances than those fed on water or sucrose (p < 0.03).

The overall flight activity of blood-fed mosquitoes was stable and continuous over the 24 h period for all diets, with an increase in activity in the latter half of the scotophase for females fed on water as well as those fed on fresh and 24 h aged urine (Fig. 3). While the urine diet significantly affected the average distance flown by blood-fed females (F_(5, 138)_ = 4.83, p = 0.0004), urea diets had no discernible effect (F_(5, 138)_ = 1.36, p = 0.24). Only blood-fed females fed on 24 h aged urine displayed an increased average flight distance compared to the other urine and control diets (fresh, p = 0.0091; 72 h, p = 0.0022; 168 h, p = 0.001; sucrose, p = 0.0017; dH_2_O, p = 0.036).

### Urine and urea affect reproductive parameters

The effect of urine and urea feeding on reproductive parameters were assessed in an oviposition bioassay (Fig. 4A), and investigated in terms of the number of eggs laid per female, as well as the size of the eggs and the newly hatched first instar larvae. The number of eggs laid by *An. arabiensis* females fed on urine varied with diet (F_(5, 222)_ = 4.38, p = 0.0008; Fig. 4B). Females fed on 24 h aged urine, post-blood meal, laid significantly more eggs than when fed on other urine diets, and similar to that laid by those fed on sucrose (Fig. 4B). Similarly, the size of eggs laid by females fed on urine differed based on diet (F_(5, 209)_ = 12.85, p < 0.0001), with females fed on 24 h aged urine and sucrose laying significantly larger eggs than those fed on water, while eggs from females fed on 168 h aged urine were significantly smaller (Fig. 4C). Moreover, the urine diets significantly affected larval size (F_(5, 187)_ = 7.86, p < 0.0001), with significantly larger larvae emerging from eggs laid by females that fed on 24 h and 72 h aged urine than those from the eggs of water-fed and 168 h aged urine-fed females (Fig. 4D).

**Figure 4.**
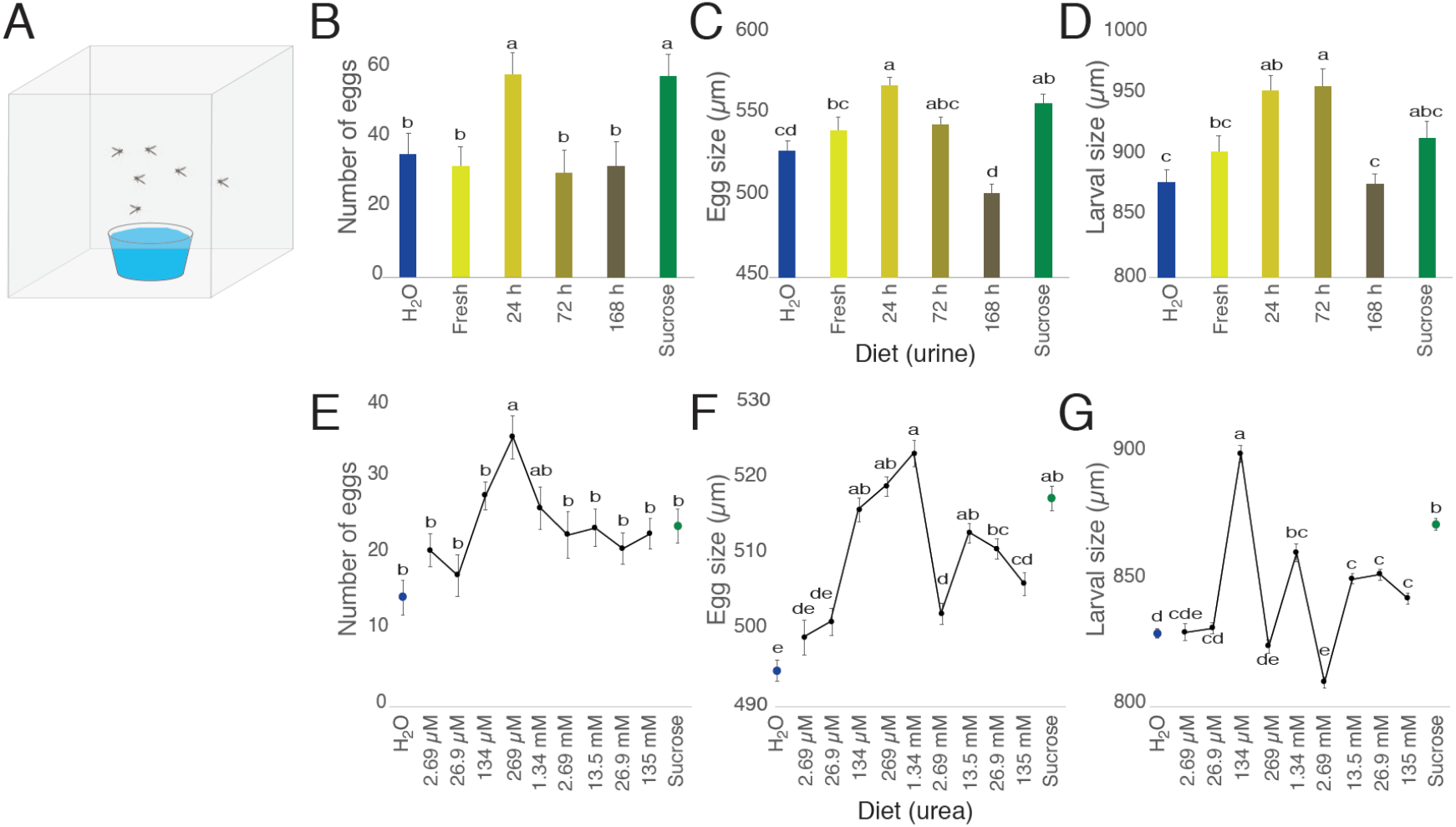
Reproductive performance of female *Anopheles arabiensis* fed on cattle urine and urea. Blood-fed female mosquitoes fed on diets consisting of fresh and aged cattle urine, various concentrations of urea, sucrose (10 %) and distilled water (H_2_O) over a period of 48 h, and then placed in a bioassay with access to an oviposition substrate for 48 h (A). The number of eggs (B, E), size of eggs (C, F) and size of larvae (D, G) were significantly affected by the diet provided (cattle urine: B-D; urea: E-G). The mean for each parameter measured with different letter designations are significantly different from one another (one-way analysis of variance with a Tukey’s *post hoc* analysis; p < 0.05). Error bars represent the standard error of the mean.

As the primary nitrogenous component of urine, urea, when offered as a diet to blood-fed females, differentially and significantly affected all of the reproductive parameters studied. The number of eggs laid by females fed on urea, post-blood meal, differed depending on the concentration of urea (F_(11, 360)_ = 4.69; p < 0.0001), with females fed on urea concentrations between 134 μM and 1.34 mM laying more eggs (Fig. 4E). Females fed on concentrations of urea at or above 134 μM laid larger eggs than those fed on water (F_(10, 4245)_ = 36.7; p < 0.0001; Fig. 4F), whereas larval size, while affected by similar concentrations of urea imbibed by the mother (F_(10, 3305)_ = 37.9; p < 0.0001), was more variable (Fig. 4G).

### Attraction of *Anopheles arabiensis* to cattle urine odour

The overall attraction to the headspace volatile extracts of cattle urine of host-seeking *An. arabiensis*, as assessed in a glass tube olfactometer (Fig. 5A), was significantly affected by the age of the urine (χ^2^ = 15.9, *df* = 4, p = 0.0032; Fig. 5B). *Post hoc* analysis revealed that 24 h aged urine odour elicited a significantly higher level of attraction compared to all other treatments (72 h: p *=* 0.0060, 168 h: p = 0.012, pentane: p = 0.00070), except fresh urine odour (p = 0.13; Fig. 5B). While there was no significant difference in the overall attraction to urine odour by blood-fed mosquitoes (χ^2^ = 8.78, *df* = 4, p = 0.067; Fig. 5C), these females were found to be significantly more attracted to the headspace volatile extract of 72 h aged urine compared to the control (p = 0.0066; Fig. 5C).

**Figure 5.**
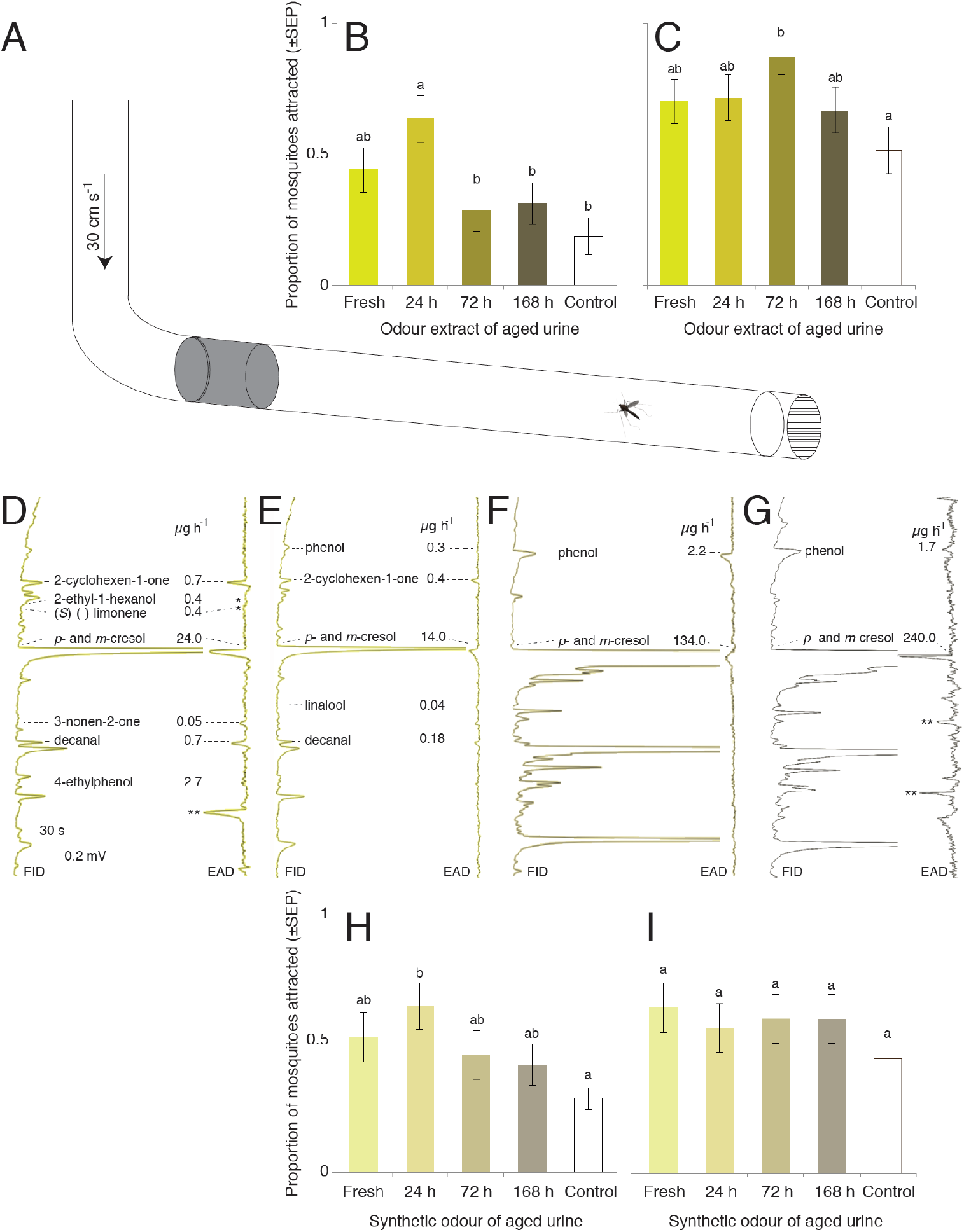
Behavioural response of host-seeking and blood-fed *Anopheles arabiensis* to natural and synthetic cattle urine odour. Diagram of the glass tube olfactometer (A). Attraction to the headspace volatile extracts of fresh and aged cattle urine of host-seeking (B) and bloodfed (C) mosquitoes. The antennal responses of host-seeking *An. arabiensis* to fractioned head-space extracts from fresh (D), 24 h (E), 72 h (F) and 168 h (G) aged cattle urine are shown. Electroantennographic detection (EAD) traces show voltage changes in response to the bioactive compounds in the headspace eluting from the gas chromatograph and detected by the flame ionization detector (FID). Scale bar indicates the amplitude of response (mV) versus the retention time (s). The identity and release rate (μg h^−1^) of the bioactive compounds are indicated. A single asterisk (*) indicates consistent low amplitude responses. Double asterisks (**) indicate irreproducible responses. Host-seeking (H) and blood-fed (I) *An. arabiensis* are differentially attracted to the synthetic blends of fresh and aged cattle urine odour. The mean proportion of mosquitoes attracted with different letter designations are significantly different from one another (one-way analysis of variance with a Tukey’s *post hoc* analysis; p < 0.05). Error bars indicate the standard error of the proportion.

### Cattle urine odour does not affect egg laying

Female *An. arabiensis*, 72 h and 120 h post-blood meal, did not demonstrate a preference for the headspace volatile extracts of fresh and aged cattle urine over that of the pentane control during oviposition (χ^2^ = 3.07, p > 0.05; Supplementary Fig. 1).

### Age affects the bioactive compounds present in cattle urine odour

For female *An. arabiensis*, the GC-EAD and GC-MS analyses identified eight, six, three and three bioactive compounds in the headspace volatile extracts of fresh, 24 h, 72 h and 168 h aged cattle urine, respectively (Fig. 5D-G). Despite the observed difference in the number of compounds eliciting an electrophysiological response, the majority of these compounds were present in each of the headspace volatile extracts collected from fresh and aged urine. Thus, only compounds that produced above-threshold physiological responses from the female antennae, for each extract, were included in further analyses.

The total volatile release rate of the bioactive compounds in the headspace collections increased from 29 μg h^−1^ in fresh urine to 242 μg h^−1^ in 168 h aged urine, predominantly due to the increase of *p*- and *m*-cresol, as well as phenol. In contrast, the release rate of other compounds, for example, 2-cyclohexen-1-one and decanal, decreased with an increasing age of the urine, correlating with the observed decrease in signal intensity (abundance) in the chromatogram (Fig. 5D-G left panel) and in the physiological response to these compounds (Fig. 5D-G right panel).

### Synthetic urine odour attracts female mosquitoes

Overall, synthetic blends approximating the natural ratio of bioactive compounds identified in the headspace volatile extracts of fresh and aged urine (Fig. 5D-G), did not appear to elicit significant attraction in host-seeking (χ^2^ = 8.15, *df* = 4, p = 0.083; Fig. 5H) or in blood-fed mosquitoes (χ^2^ = 4.91, *df* = 4, p = 0.30; Fig. 5I). However, a *post hoc* pairwise comparison among the treatments revealed a significant attraction of host-seeking mosquitoes to the synthetic blend of 24 h aged urine, as compared to the pentane control (p = 0.0086; Fig. 5H).

To assess the role of individual components in the synthetic blend of 24 h aged urine, six subtractive blends, from which individual compounds were removed, were evaluated against the full blend in a Y-tube assay. For host-seeking mosquitoes, subtraction of individual compounds from the full blend had a significant effect on the behavioural response (χ^2^ = 19.63, *df* = 6, p = 0.0032; Supplementary Fig. 2A), with all subtractive blends being less attractive than the full blend. In contrast, the removal of individual compounds from the full synthetic blend did not affect the behavioural response of blood-fed mosquitoes (χ^2^ = 11.38, *df* = 6, p = 0.077), with the exception of decanal, which resulted in a reduced level of attraction compared with the full blend (p = 0.022; Supplementary Fig. 2B).

### Synthetic cattle urine odour attracts mosquitoes under field conditions

The efficacy of the synthetic blend of 24 h aged cattle urine to attract mosquitoes under field conditions was evaluated over ten nights in a malaria endemic rural village in Ethiopia (Fig. 6A). A total of 4861 mosquitoes were captured and identified, of which 45.7 % were *An. gambiae s.l.*, 18.9 % were *An. pharoensis* and 35.4 % were *Culex spp* (Supplementary Table 1)*. Anopheles arabiensis* was the only member of the *An. gambiae* species complex to be identified by PCR analysis. On average, 320 mosquitoes were caught per night, during which time the traps baited with the synthetic blend caught more mosquitoes than the paired traps without the blend (χ^2^_(0, 3196)_ = 170.0, p < 0.0001). During each of the five control nights at the beginning, middle and end of the trial, non-baited traps were set. Similar numbers of mosquitoes were caught in each paired trap, demonstrating that there was no bias between houses (χ^2^_(0, 1665)_ = 9×10^−13^, p > 0.05) with no decline in the population over the study period. There were significantly higher numbers of mosquitoes caught in traps containing the synthetic blend compared to the control traps: host-seeking (χ^2^_(0, 2107)_ = 138.7, p < 0.0001), recently blood fed (χ^2^_(0, 650)_ = 32.2, p < 0.0001) and gravid (χ^2^_(0, 228)_ = 6.27, p = 0.0123; Supplementary Table 1). This was also reflected in the total number of mosquitoes caught: host-seeking > blood-fed > gravid > semi-gravid > males.

**Figure 6.**
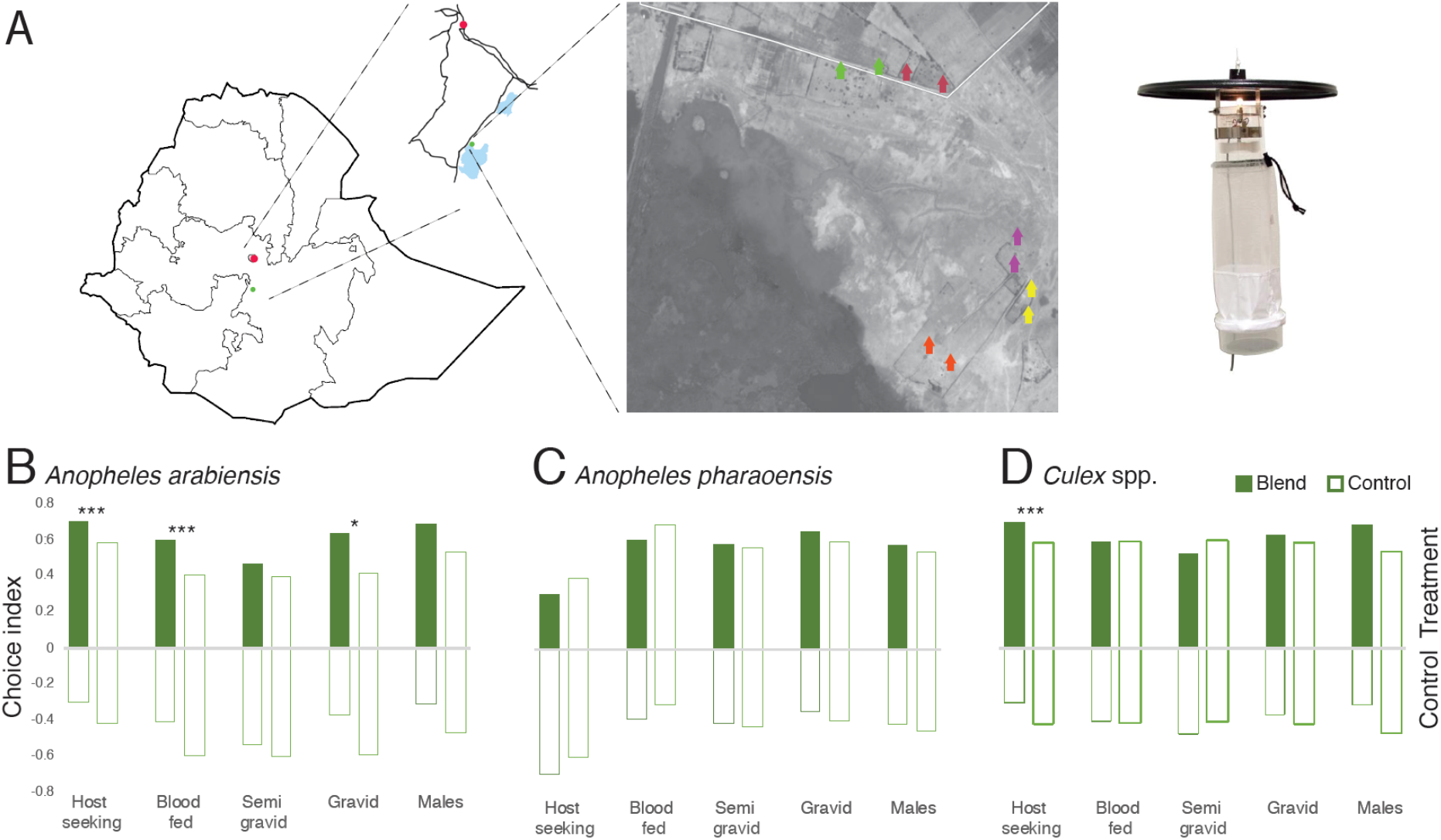
Field evaluation of the efficacy of the 24 h synthetic cattle urine odour blend. The field trials were carried out in south central Ethiopia (map), nearby the town of Meki (insert), using Centers of Disease Control (CDC) light traps (right) in paired houses using a Latin square design (aerial map) (A). CDC light traps baited with the synthetic odour differentially attracted and captured female *Anopheles arabiensis* (B), but not *Anopheles pharoensis* (C), an affect that was dependent on physiological state. In addition, the traps caught significantly higher numbers of host-seeking *Culex spp*. (D) compared to the controls. Bars on the left represent the average choice indices of mosquitoes caught in paired odour baited (green) and control (open) traps (N = 10), whereas bars on the right represent the choice indices of mosquitoes caught in paired control traps (open; N = 5). Asterisks denote the level of statistical significance (*p = 0.01 and ***p < 0.0001).

The three species were differentially caught in the traps containing the synthetic blend. A significantly higher number of host-seeking (χ^2^_(1, 1345)_ = 71.7, p < 0.0001), blood-fed (χ^2^_(1, 517)_ = 16.7, p < 0.0001) and gravid (χ^2^_(1, 180)_ = 6.11, p = 0.0134) *An. arabiensis* were caught in traps releasing the synthetic blend (Fig. 6B), whereas no difference in the number of *An. pharoensis*, at different physiological states, was found (Fig. 6C). For the *Culex spp*., only the number of host-seeking mosquitoes was found to be significantly higher in the traps baited with the synthetic blend (χ^2^_(1, 1319)_ = 12.6, p = 0.0004; Fig. 6D), compared to the control trap.

### Cattle urine odour is not a host habitat cue for mosquitoes

Host decoy traps, situated away from potential hosts between the breeding site and a rural village community in Ethiopia, were used to assess whether malaria mosquitoes use cattle urine odour as a host habitat cue. In absence of the host cue, heat, no mosquitoes were caught, with or without the presence of cattle urine odour (Supplementary Fig. 3). However, in the presence of both heat and cattle urine odour, female malaria mosquitoes were attracted and caught, although in low numbers, irrespective of the age of the urine (χ^2^_(5, 25)_ = 2.29, p = 0.13; Supplementary Fig. 3). In contrast, the water control caught no malaria mosquitoes in the presence of heat (Supplementary Fig. 3).

## Discussion

Malaria mosquitoes acquire and allocate nitrogenous compounds through compensatory feeding on cattle urine, *i.e.*, puddling, to enhance life-history traits, similar to that of other insects (Bodri, 2018; Honda et al., 2012; Molleman, 2010; Petit et al., 2019; Shen et al., 2009). Female mosquitoes locate this resource through olfaction, and are able to regulate the uptake of nitrogenous compounds in urine, including the main nitrogenous constituent of urine, urea (Dijkstra et al., 2013; Kilande et al., 2016). Depending on the life stage of the female mosquito, the nutrients within urine are allocated to enhance flight activity and survival in host-seeking females, and survival and reproductive traits in blood-fed individuals. As such, urine puddling plays an important nutritive role for malaria vectors that eclose as undernourished adults (Van Handel, 1965). This finding has significant epidemiological consequences, as females increase their life expectancy, activity and reproductive output, all of which affect vectorial capacity.

The VOC profile of urine changes with age as a result of microbial activity (Kilande et al., 2016; Okech and Hassanali, 1990; Storer et al., 2011; Troccaz et al., 2013). Host-seeking female *An. arabiensis* are attracted to the VOCs of fresh and 24 h aged urine (this study, Kweka et al., 2009; Mahande et al., 2010), which is different from that found for other dipterans, including tsetse and tabanids, which prefer VOCs of older aged urine (Mihok and Mulye, 2010; Okech and Hassanali, 1990; Vale et al., 1988). The overall complexity of VOCs increases as the urine ages, with phenol and phenolic derivatives as the predominant VOCs (this study, Okech and Hassanali, 1990; Baldacchino et al., 2013). While blends of phenolic VOCs are sufficient to elicit attraction in tsetse and tabanids (Baldacchino et al., 2013; Madubunyi et al., 1996; Mihok and Mulye, 2010; Vale et al., 1988), these fail to do so in *An. arabiensis*, as corroborated by (Mahande et al., 2010) and (Kweka et al., 2009). In contrast, the blends of antennally-detected VOCs that attract female *An. arabiensis* are more complex. These blends also contain phenol, *p*- and *m*-cresol, although at lower release rates compared to that found in the older urine. A synthetic blend of these phenolic compounds, along with three addition antennally active VOCs is required to recapitulate the behavioural response of host-seeking females to cattle urine under laboratory conditions. This suggests an evolutionarily conserved function of phenolic compounds among dipterans, however the context in which these phenolics are presented is adaptive for different species. When assessed under field conditions, the same blend elicited attraction of host-seeking *An. arabiensis* and *Culex spp*. females, but not of *An. pharoensis*, emphasising a conserved, yet species-dependent, response. Cattle urine VOCs have been proposed to act as host habitat cues, *i.e.*, long-range attractants that indicate the presence of a potential host within a particular area, for tsetse, tabanids and other non-Culicidae flies (Webster and Cardé, 2017). Mosquitoes, however, do not appear to use cattle urine as a host habitat cue, emphasising a different ecological function in Culicidae (mosquitoes) and non-Culicidae flies.

While the synthetic odour of 24 h aged urine attracted recently blood-fed and gravid *An. arabiensis* in the field, this was not observed under laboratory conditions. In contrast, blood fed females exhibited a strong attraction to the background humid air, with little to no effect of the cattle urine VOCs. These behavioural results in the laboratory are likely confounded by the fact that humidity itself is a strong preoviposition attractant (Okal et al., 2013), but is a prerequisite for the bioassay. Cattle urine has been proposed to act as an oviposition attractant for gravid mosquitoes (Kweka et al., 2011), however this was not supported in this study, leading us to search for other plausible explanations for the attraction of mosquitoes to cattle urine.

Fresh urine mainly contains salts and nitrogenous compounds, two nutrient classes frequently sought for by insects using supplemental feeding to increase fitness (Molleman, 2010). Host-seeking and blood-fed *An. arabiensis* actively imbibe cattle urine, at a similar level as water intake, irrespective of the age of the urine, which may be due to similar overall levels of nitrogen in fresh and aged urine. As cattle urine ages, microbes make use of the nitrogenous compounds in urine, particularly hydrolysing urea to ammonia, resulting in a changing complexity of the microbial communities (Kilande et al., 2016). While females do not display any obvious feeding preference for fresh or aged urine, mosquitoes demonstrate a dose-dependent response to urea, revealing that both host-seeking and blood-fed mosquitoes regulate their intake of nitrogenous compounds. Host-seeking mosquitoes imbibe a wide range of urea concentrations. However, these females display optimal intake volumes of urea at concentrations similar to those present in fresh and 24 h aged cattle urine (Dijkstra et al., 2013; Kilande et al., 2016), and which does not differ from the sucrose control. While blood-fed mosquitoes imbibe lower volumes of urea and water than host-seeking females, as recently blood-fed females are constrained by the previous meal, these females display a lower threshold of response to urea. Mosquitoes are unable to metabolise urea, and likely use gut bacteria that possess ureases to hydrolyse urea to ammonia (Chen et al., 2017; Kämpfer et al., 2011). Midgut tissues and fat bodies in mosquitoes are able to convert ammonia into the amino acids, glutamate, glutamine, alanine and proline (Scaraffia et al., 2005; Scaraffia et al., 2010), which are important components of yolk proteins and, in the case of proline, can be used as an energy source for flight (Scaraffia and Wells, 2003).

Allocation patterns of assimilated nitrogenous nutrients from urine, including urea, are not independent, as these nutrients are allocated as a function of physiological state to survival, flight and reproduction in order to provide the life history traits demonstrated in host-seeking and blood-fed mosquitoes. This conforms with the general aspect of nutrient allocation found for other insects, in which life history traits are constrained by one or more limiting nutrients (Boggs, 2009; Raubenheimer et al., 2009). The need to allocate nutrients to more than one trait at the same time can generate physiological trade-offs among those traits (Boggs, 2009), as demonstrated in a pair-wise manner for host-seeking (flight vs. survival) and blood-fed (survival vs. reproduction) mosquitoes. The need to increase acquisition of a food type containing a limiting nutrient has been shown to result in excess consumption of nutrients that are deleterious to another allocation target (Boggs, 2009; Lee et al., 2008). Such trade-offs may explain why host-seeking female mosquitoes are attracted to and imbibe 72 h aged urine, which contains toxic microbiota (Kilande, Tenywa, Rwakaikara-Silver, et al., 2016), increasing the distance flown but resulting in a significantly reduced life span. On the other hand, the acquisition, by blood-fed females, of excess nutrients in 24 h aged urine, allow allocation of these resources to more than one trait, *i.e.*, survival, flight and reproduction. This demonstrates that compensatory feeding on urine can be used for similar purposes as multiple blood meals within one gonotrophic cycle (Briegel and Horler, 1993). This allocation framework (Boggs, 2009; Raubenheimer et al., 2009) provides a mechanistic understanding of life history patterns and how resources are allocated to survival, dispersal and reproduction.

Compensatory feeding for nitrogenous compounds, in the form of multiple blood meals, has been shown in *An. gambiae s. l.* to either be required for egg development in females with low teneral reserves, or to enhance the number and condition of eggs developing in a single gonotrophic cycle (Beier, 1996; Gillies, 1954; Scott and Takken, 2012; Takken et al., 1998). However, blood feeding is risky and presents a trade-off for the female between reproduction and survival (Anderson and Roitberg, 1999) making another, low risk, source for nitrogenous compounds, *e.g*., cattle urine, an adaptive alternative. The increased survival, as well increased numbers and sizes of eggs laid following a compensatory urine or urea meal by a blood fed female reflects that which is observed following multiple blood meals (Gillies, 1954; Scott and Takken, 2012; Takken et al., 1998). This suggests that *An. arabiensis* may minimise the trade-off between the need for nitrogen resources to enhance reproductive traits and survival by making use cattle urine.

Blood-fed mosquitoes allocate the bulk of the nitrogenous compounds from compensatory feeding to reproduction, and in part to survival, while host-seeking mosquitoes predominantly use these to fuel flight (Gaviraghi et al., 2019), analogous to that described for other insects (Teulier et al., 2016; Tigreros and Davidowitz, 2019). The immediate and sustained increase in flight activity by host-seeking females following a urea meal suggests that this resource can be used directly to fuel flight activity, potentially through the use of the combination of the previously described conversion of ammonia to proline (Scaraffia et al., 2010) and its further oxidation of proline to fuel flight muscles (Gaviraghi et al., 2019; Scaraffia and Wells, 2003). The dawn activity pattern demonstrated by host-seeking females fed on fresh urine reflects that observed for *An. gambiae* females engaging in compensatory blood feeding (Klowden and Briegel, 1994). Host-seeking females fed on 24 h and 168 h aged urine, on the other hand, demonstrate an abnormal activity throughout photophase, suggesting that these females may be nutrient seeking during this time. Feeding on 72 h aged urine resulted in activity patterns similar to those demonstrated post-urea feeding, reflecting the high levels of ammonia present in the diet at this time, as a result of microbial activity (Kilande et al., 2016). Thus, mosquitoes have the capacity to use cattle urine, and its main nitrogenous component urea, as fuel for flight.

Malaria mosquitoes demonstrate complex behavioural and physiological strategies to adapt to their environment. Feeding on cattle urine compensates for the need to take multiple blood meals by enhancing life history traits in a state dependent manner. This is likely to affect vectorial capacity by increasing the probability of daily survival and vector density, while decreasing the interaction between the vector and the host, by reducing the need for multiple blood meals. Urine meals provide an alternate, non-blood, nitrogen source and reduce the number of undernourished females requiring a pre-blood meal for metabolic energy prior to egg development. As such, compensatory feeding on cattle urine, or other nitrogen-rich resources, should be taken into consideration in future models of vectorial capacity. To this end, further studies are required to establish the natural role of cattle urine feeding in malaria mosquitoes, and how this behaviour may be manipulated in future vector management strategies.

## Materials and Methods

### Mosquito rearing

*Anopheles arabiensis* (Dongola strain) were maintained at 25 ± 2 °C, 65 ± 5 % RH and at a 12:12 h light: dark cycle. Larvae were reared in plastic trays (20 cm × 18 cm × 7 cm), filled with distilled water, and fed on Tetramin® fish food (Tetra Werke, Melle, DE). Pupae were collected in 30 ml cups (Nolato Hertila, Åstorp, SE) and transferred to Bugdorm cages (30 cm × 30 cm × 30 cm; MegaView Science, Taichung, TW) for the adults to emerge. Adults were provided with 10 % sucrose solution *ad libitum* until 4 days post-emergence (dpe), at which time host-seeking females were either provided with the diet immediately, or starved overnight with access to distilled water, prior to experiments, as described below. Females used for the flight tube experiments were only starved for 4-6 h with *ad libitum* access to water. To prepare blood-fed mosquitoes for subsequent bioassays, 4 dpe females were provided defibrinated sheep blood (Håtunalab, Bro, SE) using a membrane feeding system (Hemotek Discovery Workshops, Accrington, UK). Fully engorged females were subsequently transferred to separate cages and provided either a diet directly, as described below, or *ad libitum* access to 10 % sucrose for 3 days, prior to the experiments described below. The latter females were used for the flight tube bioassays and were transferred to the experimental room, then starved with *ad libitum* access to distilled water 4-6 h prior to the experiments.

### Quantification of urine and urea imbibed

Feeding assays were used to quantify the consumption of urine and urea by adult *An. arabiensis* females. Host-seeking and blood-fed females were provided with diets containing a 1 % dilution of fresh and aged cattle urine, various concentrations of urea, as well as two controls, 10 %sucrose and water, for 48 h. In addition, a food colourant (1 mg ml^−1^ xylene cyanole FF; CAS 2650-17-1; Sigma-Aldrich, Stockholm, SE), was added to the diet and provided in a 4 × 4 matrix of 250 μl microfuge tubes (Axygen Scientific, Union City, CA, US; Fig. 1A) filled to the rim (ca. 300 μl). To avoid competition among mosquitoes and the potential influence of the colour of the dye, ten mosquitoes were placed in large Petri dishes (12 cm diameter, 6 cm height; Semadeni, Ostermundigen, CH; Fig. 1A) in complete darkness at 25 ± 2 °C and 65 ± 5 % RH. These experiments were replicated from 5 to 10 times. Following exposure to the diets, the mosquitoes were placed at −20 °C until further analysis.

To release the diet imbibed, mosquitoes were placed individually in 1.5 ml microfuge tubes containing 230 μl of distilled water, and the tissues disrupted using a disposable pestle and cordless motor (VWR International, Lund, SE), and then centrifuged at 10 krpm for 10 min. The supernatants (200 μl) were transferred to a 96-well microplate (Sigma-Aldrich) and the absorbance (λ620 nm) determined using a spectrophotometer-based microplate reader (SPECTROStar^®^ Nano, BMG Labtech, Ortenberg, DE). Alternatively, the mosquitoes were ground in 1 ml of distilled water, 900 μl of which was transferred to a cuvette for spectrophotometric analysis (λ620 nm; UV 1800, Shimadzu, Kista, SE). To quantify the diet imbibed, a standard curve was prepared by a serial dilution resulting in a range of 0.2 μl to 2.4 μl of 1 mg ml^−1^ xylene cyanol. Then, the optical density of known dye concentrations was used to determine the volume of diet imbibed by each mosquito.

The volumetric data were analysed using a one-way analysis of variance (ANOVA) followed by a Tukey’s *post hoc* pairwise comparison (JMP Pro, v14.0.0, SAS Institute Inc., Cary, NC, US, 1989-2007). Linear regression analysis described the concentration dependent urea intake and comparisons were made between the responses of host-seeking and blood-fed mosquitoes (GraphPad Prism v8.0.0 for Mac, GraphPad Software, San Diego, CA, US).

### Urine and nitrogen analysis

Approximately 20 μl of sample urine from each age category was bound on Chromosorb^®^ W/AW (10 mg 80/100 mesh, Sigma Aldrich), and enclosed in tin capsules (8 mm × 5 mm). The capsule was inserted into a combustion chamber of a CHNS/O analyser (Flash 2000, Thermo Fisher Scientific, Waltham, MA, US) to determine the nitrogen content of fresh and aging urine, according to the manufacturer’s protocol. Total nitrogen (g N l^−1^) was quantified based on known concentrations of urea used as the standard.

### Survival analysis

To assess the effect of diet on the survival of host-seeking and blood-fed females, mosquitoes were placed individually into large Petri dishes (diameter 12 cm, height 6 cm; Semadeni), with a mesh covered hole in the lid (3 cm diameter) for ventilation and diet provision. The diets, consisting of a 1 % dilution of fresh and aged cattle urine, four concentrations of urea, as well as two controls, 10 % sucrose and water, were provided directly after 4 dpe. Each diet was pipetted onto dental cotton rolls (DAB Dental AB, Upplands Väsby, SE) inserted into 5 ml syringes (Thermo Fisher Scientific, Gothenburg, SE), with the plunger removed, and then placed on top of the Petri dishes (Fig. 1A). The diets were replaced daily. The experimental room was maintained as described above. Surviving mosquitoes were counted twice daily, while discarding dead mosquitoes, until the final mosquito died (n = 40 per treatment). The survival of the mosquitoes feeding on the respective diets was analysed using Kaplan-Meyer survival curves and log rank test statistics for survival distribution comparison between diets (IBM SPSS Statistics 24.0.0.0).

### Tethered flight assay

A custom-made mosquito flight-mill, based on (Attisano et al., 2015), was made from 5 mm thick clear acrylic panels (10 cm W × 10 cm L × 10 cm H) lacking front and back panels (Fig. 3: top). A pivot assembly, with a vertical tube constructed from a gas chromatography column (0.25 mm i.d; 7.5 cm L) glued to insect pins at both ends, was suspended between a pair of neodymium magnets, 9 cm apart. A horizontal tube made of the same material (6.5 cm L) bisected the vertical tube and created a tethering arm and an arm that carried a small piece of aluminium foil as a photo interruption signal.

Prior to tethering, 24 h starved females, were provided access to the diets described above for 30 min. Fully fed female mosquitoes were then individually anaesthetized on ice for 2-3 min and glued onto an insect pin using bee’s wax (Joel Svenssons Vaxfabrik AB, Munka Ljungby, SE) on their mesothorax, and then tethered onto the arm of the horizontal tube of the flight-mill. Each flight revolution was logged by a customized data logger, then stored and displayed using the PC-Lab 2000™ software (v4.01; Velleman, Gavere, BE). The flight mill was placed in a climate conditioned room (12 h: 12 h, light: dark, 25 ± 2 °C, 65 ± 5 % RH).

To visualise the pattern of flight activity, the overall distance flown (m) and the overall number of bouts of continuous flight activity were calculated each hour over the course of a 24 h period. In addition, the average distance flown by an individual female was compared among the various treatments and analysed using a one-way analysis of variance followed by a Tukey *post hoc* analysis (JMP Pro, v14.0.0, SAS Institute Inc.), in which the average distance was considered a dependent variable, while the treatments were the independent factors. Moreover, the average number of bouts was also calculated in 10-min increments.

### Reproductive performance

To assess the effect of diet on the reproductive performance of *An. arabiensis*, six females (4 dpe) were transferred to Bugdorm cages (30 cm × 30 cm × 30 cm) directly after blood feeding, and then provided with experimental diets, as described above, for 48 h. Diets were then removed and oviposition cups (30 ml; Nolato Hertila), filled with 20 ml distilled water, were provided on the third day and made available for 48 h, replacing the cups every 24 h. Each diet regime was replicated 20-50 times. The eggs were counted and recorded for each experimental cage. A subsample of eggs was used to assess the average size and variation among the lengths of individual eggs (n ≥ 200 per diet) using a Dialux-20 microscope (DM1000; Ernst Leitz Wetzlar, Wetzlar, DE) equipped with a Leica camera (DFC 320 R2; Leica Microsystem Ltd, DE). The remaining eggs were maintained in a climate-controlled chamber under standard rearing conditions for 24 h, and a subsample of recently emerged 1^st^ instar larvae (n ≥ 200 per diet) were measured, as above. The number of eggs, as well as the size of both eggs and larvae, were compared among the various treatments and analysed using a one-way analysis of variance followed by a Tukey *post hoc* analysis (JMP Pro, v14.0.0, SAS Institute Inc.).

### Headspace volatile collections from fresh and aged cattle urine

Headspace volatiles from fresh (1 h post-sampling), 24 h, 72 h and 168 h aged urine were collected from samples collected from Zebu cattle, Arsi race. For convenience and availability, the urine sample collections were carried out early in the morning while the cattle were still in the shed. Urine samples were collected from ten individuals, with 100-200 ml of each sample transferred into separate polyamide roasting bags (Toppits Cofresco, Frischhalteprodukte GmbH and Co., Minden, DE) placed inside a 3 l polyvinylchloride plastic bucket with a lid. Headspace volatiles from each individual cattle urine sample were either collected directly (fresh) or following maturation for 24 h, 72 h and 168 h at room temperature, *i.e*., each urine sample was represented in each of the age groups.

For the headspace volatile collection, a closed loop system was used, by circulating an activated charcoal-filtered airstream (100 ml min^−1^) through the polyamide bag onto an adsorbent column, using a diaphragm vacuum pump (KNF Neuberger, Freiburg, DE), for 2.5 h. As a control, headspace collection from an empty polyamide bag was performed. The adsorbent column was made of Teflon tubing (5.5 cm × 3 mm i.d.) holding 35 mg Porapak Q (50/80 mesh; Waters Associates, Milford, MA, US) between glass wool plugs. The columns were rinsed with 1 ml re-distilled *n*-hexane (Merck, Darmstadt, DE) and 1 ml pentane (99.0 % pure solvent GC grade, Sigma Aldrich) before use. Adsorbed volatiles were eluted with 400 μl pentane. Headspace collections were pooled and then stored at −20 °C until used for further analyses.

### Attraction of *Anopheles arabiensis* to fresh and aged cattle urine odour

Behavioural responses of host-seeking and blood-fed *An. arabiensis* mosquitoes to the head-space volatile extracts collected from fresh, 24 h, 72 h and 168 h aged urine were analysed using a straight glass tube olfactometer (Majeed et al., 2014). The experiments were conducted during the peak host-seeking activity period, ZT 13-15, of *An. arabiensis* (Jones et al., 1967). The glass tube olfactometer (80 cm × 9.5 cm i.d.) was illuminated with red light from above at 3 ± 1 lx. A charcoal-filtered and humidified air stream (25 ± 2 °C, 65 ± 2 % relative humidity) passed through the bioassay at 30 cm s^−1^. The air passed through a series of stainless-steel mesh screens to generate a laminar flow and a homogenous plume structure. Dental cotton roll dispensers (4 cm × 1 cm; L:D; DAB Dental AB), suspended from a 5 cm wire coil at the upwind end of the olfactometer, were used and the stimulus replaced every 5 min. For the analysis, 10 μl of each headspace extract, at a 1:10 dilution, was used as the stimulus. An equivalent amount of pentane was used as a control. Individual host-seeking or blood-fed mosquitoes were placed in separate release cages 2-3 h before the onset of the experiments. The release cage was placed at the down-wind end of the olfactometer, and mosquitoes were allowed 1 min to acclimatize before the butterfly valve of the cage was opened for their release. Attraction to either treatment or control was analysed as the proportion of mosquitoes that made source contact within 5 min after release. Each headspace volatile extract and control was replicated at least 30 times, and to avoid any day effect, the same number of treatments and controls were tested on each experimental day. Responses of host-seeking and blood-fed *An. arabiensis* to the headspace collections were analysed using a nominal logistic regression followed by pairwise comparisons of the odd’s ratios (JMP Pro, v14.0.0, SAS Institute Inc.).

### Oviposition of *Anopheles arabiensis* in response to fresh and aged cattle urine odour

The oviposition response of *An. arabiensis* to the headspace extracts of fresh and aged cattle urine was analysed in Bugdorm cages (30 cm × 30 cm × 30 cm; MegaView Science). Plastic cups (30 ml; Nolato Hertila) filled with 20 ml distilled water provided the oviposition substrate, and were placed in opposite corners of the cage, 24 cm apart. The treatment cup was conditioned with 10 μl of each headspace extract, at a 1:10 dilution. An equivalent amount of pentane was used to condition the control cup. Treatment and control cups were exchanged in between each experiment to control for location effects. Ten blood-fed females were released into the experimental cages at ZT 9-11, and the number of eggs in the cups were counted after 24 h. An oviposition index was calculated by: (number of eggs laid in treatment cups – number of eggs laid in the control cups)/(total number of eggs). Each treatment was replicated 8 times.

### Combined gas chromatography and electroantennographic detection (GC-EAD)

Combined gas chromatography and electroantennographic detection (GC-EAD) analyses of female *An. arabiensis* were performed as previously described (Wondwosen et al., 2016). Briefly, an Agilent Technologies 6890 GC (Santa Clara, CA, US), equipped with an HP-5 column (30 m × 0.25 mm i.d., 0.25 μm film thickness, Agilent Technologies), was used to separate the headspace volatile extracts of fresh and aged urine. Hydrogen was used as the mobile phase at an average linear flow rate of 45 cm s^−1^. Each sample (2 μl) was injected in splitless mode, for 30 s, at an injector temperature of 225 °C. The GC oven temperature was programmed from 35 °C (3 min hold) at 10 °C min^−1^ to 300 °C (10 min hold). At the GC effluent splitter, 4 psi of nitrogen was added and split 1:1 in a Gerstel 3D/2 low dead volume four-way cross (Gerstel, Mülheim, DE) between the flame ionization detector and the EAD. The GC effluent capillary for the EAD passed through a Gerstel ODP-2 transfer line, which tracked the GC oven temperature plus 5 °C, into a glass tube (10 cm × 8 mm), where it was mixed with charcoal-filtered, humidified air (1.5 l min^−1^). The antenna was placed 0.5 cm from the outlet of this tube. Each individual mosquito accounted for a single replicate, and at least three replicates were performed for each age of the urine samples, for host-seeking mosquitoes.

### Chemical analysis

Bioactive compounds in the headspace collections of fresh and aged cattle urine, eliciting an antennal response in the GC-EAD analyses, were identified using a combined GC- and mass spectrometer (GC-MS; 6890 GC and 5975 MS; Agilent Technologies), operated in the electron impact ionization mode at 70 eV. The GC was equipped with an HP-5MS UI coated fused silica capillary column (60 m × 0.25 mm i.d., 0.25 μm film thickness), and helium was used as the mobile phase at an average linear flow rate of 35 cm s^−1^. A 2 μl sample was injected using the same injector settings and oven temperatures as for the GC-EAD analysis. Compounds were identified according to their retention times (Kovát’s indices) and mass spectra, in comparison with custom-made and NIST14 libraries (Agilent). Identified compounds were confirmed by the injection of authentic standards (Supplementary Table 2). For quantification, heptyl acetate (10 ng, 99.8 % chemical purity, Aldrich) was injected as an external standard.

### Behavioural assays with synthetic odour blends

To assess the efficacy of the synthetic odour blends, composed of the bioactive compounds identified in fresh and aged urine, to attract host-seeking and blood-fed *An. arabiensis*, the same olfactometer and protocol were used as described above. The synthetic blends mimicked the composition and ratio of compounds in the pooled headspace volatile extracts of fresh, 24 h, 48 h, 72 h and 168 h aged urine (Fig. 5D-G; Supplementary table 2). For the analysis, 10 μl of a 1:100 dilution of the full synthetic blends, at overall release rates ranging from approximately 140-2400 ng h^−1^, were used to assess attraction of host-seeking and blood-fed mosquitoes. Thereafter, subtractive blends, in which single compounds of the full blend were removed, were tested against the full blend. Responses of host-seeking and blood-fed *An. arabiensis* to the synthetic and subtractive blends were analysed using a nominal logistic regression followed by pairwise comparisons of the odd’s ratios (JMP Pro, v14.0.0, SAS Institute Inc.).

### Assessment of Cattle Urine Odour as a Host Habitat Cue

To assess whether cattle urine serves as a host habitat cue for malaria mosquitoes, fresh and aged cattle urine, collected as above, as well as water, were placed into mesh-covered 3 l buckets (100 ml), with side perforations, and set on top of host decoy traps (BG-HDT version; Bio-Gents, Regensburg, DE). The ten traps were placed 50 m apart in a pasture, separated 400 m away from a village community (Sile, Ethiopia, 5°53′24″N, 37°29′24″E) and devoid of cattle, situated between the permanent breeding site and the village. Five traps were heated to simulate the presence of a host, while five traps remained unheated. The position of each treatment was rotated nightly for a total of five nights. Comparisons among the number of mosquitoes caught in traps baited with different ages of the urine were made using logistic regression with a beta binomial distribution (JMP Pro, v14.0.0, SAS Institute Inc.).

### Field Evaluation of Synthetic Cattle Urine Odour

The efficacy of the synthetic 24 h cattle urine odour blend to attract wild mosquitoes in the field was assessed in a malaria-endemic village nearby the town of Meki in the Oromia region of Ethiopia (8°11′08″N, 38°81′70″E; Fig. 6A). The study was conducted between mid-August to mid-September prior to the annual indoor residual spraying, in conjunction with the long rainy season. Five pairs of houses (20-50 m apart), located in the periphery of the village were selected for the study (Fig. 6A). The criteria used to select the houses were: no animals were allowed to be kept inside the houses, no cooking (smoking fire wood or charcoal) was allowed indoors (at least during the trial period), and houses with a maximum of two inhabitants, sleeping under a non-insecticide treated bed nets. Ethical approval was obtained from the Institutional Research Ethics Review Board, College of Natural Sciences, (CNS-IRB), Addis Ababa University (IRB/022/2016), according to the guidelines set out by the World Medical Association Declaration of Helsinki. Consent from each household head was obtained with assistance of health extension workers. The whole process was endorsed by the local administration at district and ward (‘Kebele’) level. The experimental design followed a 2 Í 2 Latin square design, in which the synthetic blend and control were assigned to paired houses at the first night and exchanged between the houses on the next experimental night. This procedure was replicated ten times. In addition, to estimate the activity of mosquitoes in the selected houses, CDC traps, without synthetic blend dispensers, were set to operate during the same hours of the day, at the beginning, middle and end of the field trials for five nights.

The synthetic blend, containing the six bioactive compounds in their natural ratio (7:9:156:156:1:4; Fig. 5D-G; Supplementary table 2) was dissolved in heptane (97.0 % solvent GC grade, Sigma Aldrich), and released at 140 ng h^−1^ using cotton wick dispensers (Wondwosen et al., 2016). The wick dispensers allow for the release of all compounds in constant proportions throughout the 12 h experiment. Heptane was used as a control. The vials were suspended next to the entrance point of a Center for Disease Control and Prevention (CDC) light trap (John W. Hock Company, Gainesville, FL, US; Fig. 6A). The traps were suspended 0.8 – 1 m above the ground next to the foot side of a bed with a volunteer sleeping under an untreated bed net and operated between 18h00 and 06h30. Caught mosquitoes were sorted by sex and physiological state (unfed, fed, semi-gravid and gravid (WHO, 1975). Subsequently, the mosquitoes were identified morphologically to species (Gillies and Coetzee, 1987; Verrone, 1962) and placed in 1.5 ml microfuge tubes with dry silica gel. Five per cent of the mosquitoes that were morphologically identified as *An. gambiae s.l.* were subsequently screened using polymerase chain reaction (PCR) analysis to identify the member of the species complex (Wilkins et al., 2006). To assess the effect of treatment to that of the control in the field studies, trap captures of the paired houses were analysed using a nominal logistic fit model, in which attraction was the dependent variable and treatment (synthetic blend vs. control) the fixed effect (JMP^®^ 14.0.0. SAS Institute Inc.). Here, we report the χ^2^ and p-value from the Likelihood Ratio Test.

## Acknowledgements

We thank Dr Elsa Quillery for assisting in the PCR analysis of field collected mosquitoes. We also thank Yared Debebe for his assistance with the HDT experiments.

## Competing interests

The authors declare that they have no competing interest.

## Supplementary files

**Supplementary table 1:** Species, sex and gonotrophic state of the mosquitoes captured in CDC light traps baited with the synthetic odour blend of 24 h aged cattle urine or heptane control.

|  |  | Host-seeking | Blood fed | Semi-gravid | Gravid | Males |
| --- | --- | --- | --- | --- | --- | --- |
| CDC light traps | *Anopheles arabiensis* | 466 | 154 | 30 | 71 | 29 |
|  | *Anopheles pharoensis* | 141 | 25 | 14 | 23 | 5 |
|  | *Culex* spp. | 562 | 100 | 19 | 17 | 9 |
| CDC light traps baited with synthetic blend | *Anopheles arabiensis* | 879 | 363 | 85 | 109 | 37 |
|  | *Anopheles pharoensis* | 471 | 123 | 29 | 62 | 26 |
|  | *Culex* spp. | 757 | 164 | 22 | 57 | 12 |

**Supplementary table 2.** Synthetic compounds used for electrophysiological and behavioural analyses.

| Compound | Compound class | CAS No. | Purity (%) | Supplier |
| --- | --- | --- | --- | --- |
| 2-ethyl-1-hexenol | aliphatic alcohol | 104-76-7 | 99 | Sigma-Aldrich |
| decanal* | aliphatic aldehyde | 112-31-2 | 98 | Sigma-Aldrich |
| 3-nonen-2-one | aliphatic ketone | 18402-83-0 | 95 | Sigma-Aldrich |
| 2-cyclohexen-1-one* | cyclic, aliphatic ketone | 930-68-7 | 96 | VWR |
| phenol* | aromatic alcohol | 108-95-2 | 99.5 | Sigma-Aldrich |
| *m*-cresol* | aromatic alcohol | 108-39-4 | 97 | Sigma-Aldrich |
| *p*-cresol* | aromatic alcohol | 106-44-5 | 99 | Sigma-Aldrich |
| 4-ethylphenol | aromatic alcohol | 123-07-9 | 99 | Sigma-Aldrich |
| *S*-(-)-limonene | monoterpenic hydrocarbon | 5989-54-8 | 95 | Sigma-Aldrich |
| linalool* | monoterpenic alcohol | 78-70-6 | 97 | Sigma-Aldrich |

**Supplementary figure 1:**
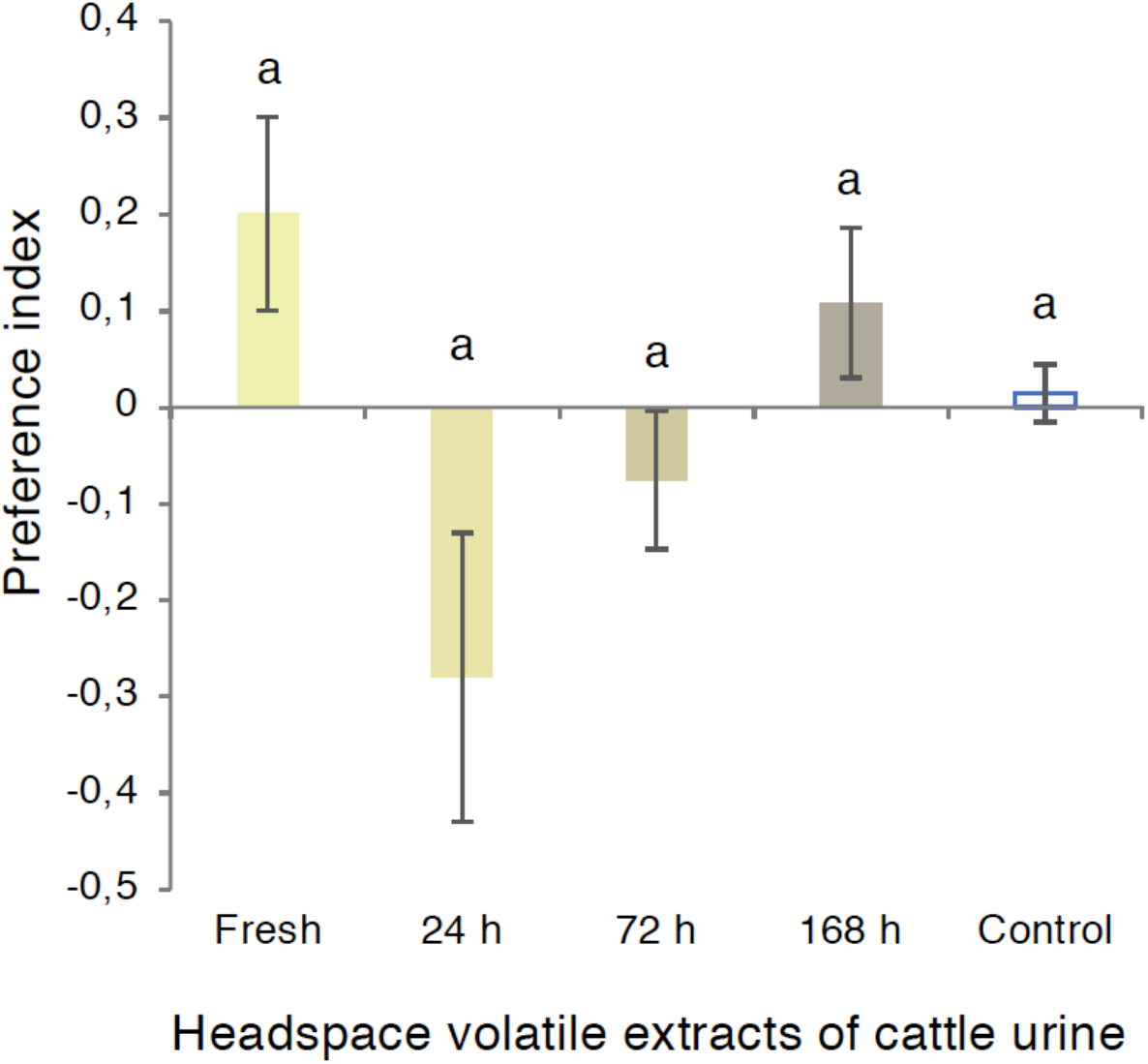
Blood-fed *Anopheles arabiensis* display no oviposition preference for the headspace volatile extracts of fresh and aged cattle urine. Letter designations indicate no significant difference from one another (one-way analysis of variance with a Tukey’s *post hoc* analysis; p > 0.05). Error bars indicate the standard error of the proportion.

**Supplementary figure 2.**
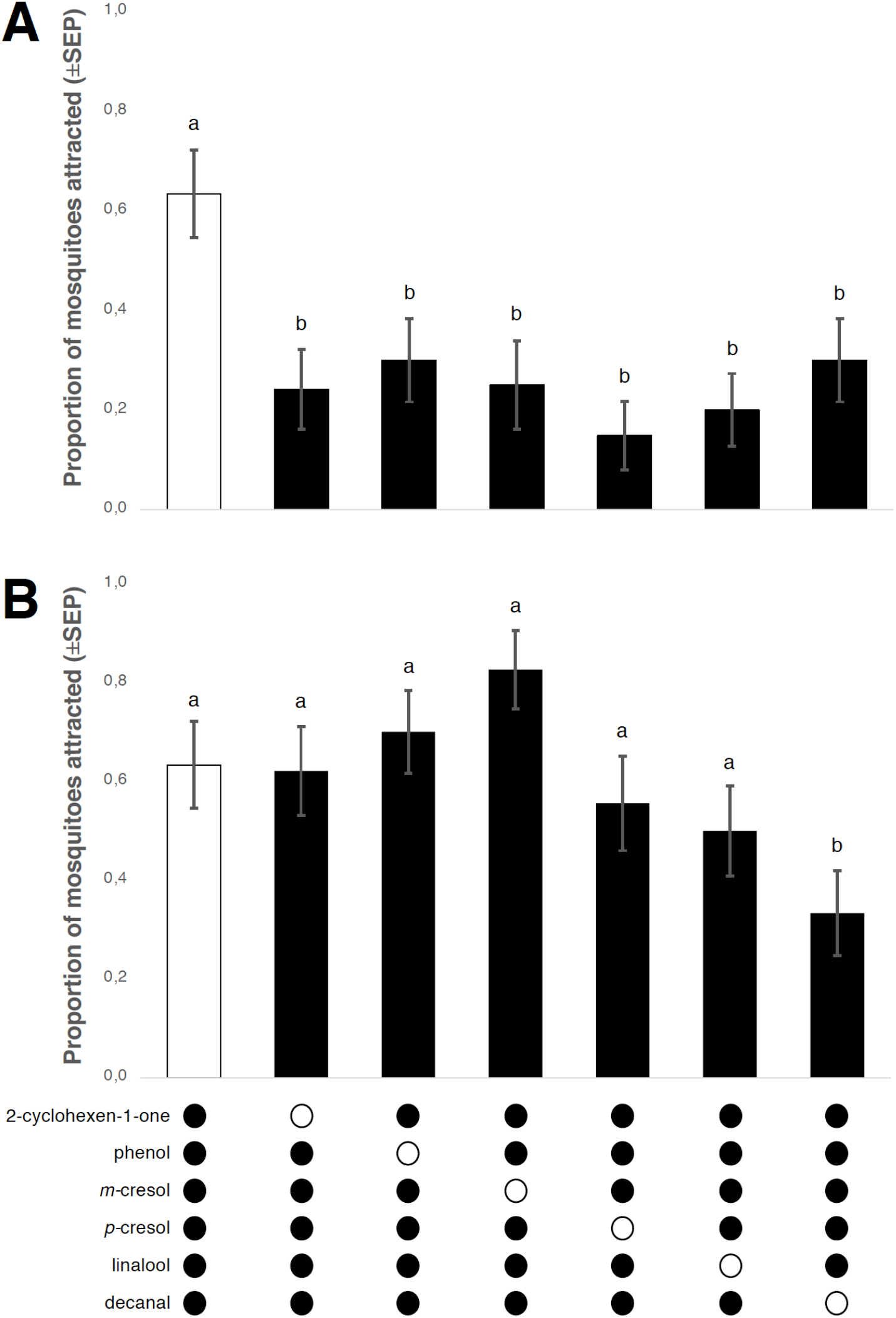
Behavioural responses of host-seeking (A) and blood-fed (B) *Anopheles arabiensis* to the full and subtractive synthetic blends of 24 h aged cattle urine. The removal of single components from the synthetic blend (open circles) differentially and significantly affected the response of the females from both physiological states. Different lowercase letters indicate significant differences as determined by a one-way analysis of variance followed by a Dunnett′s *post hoc* analysis (*p* < 0.05). Error bars represent the standard error of proportion.

**Supplementary figure 3.**
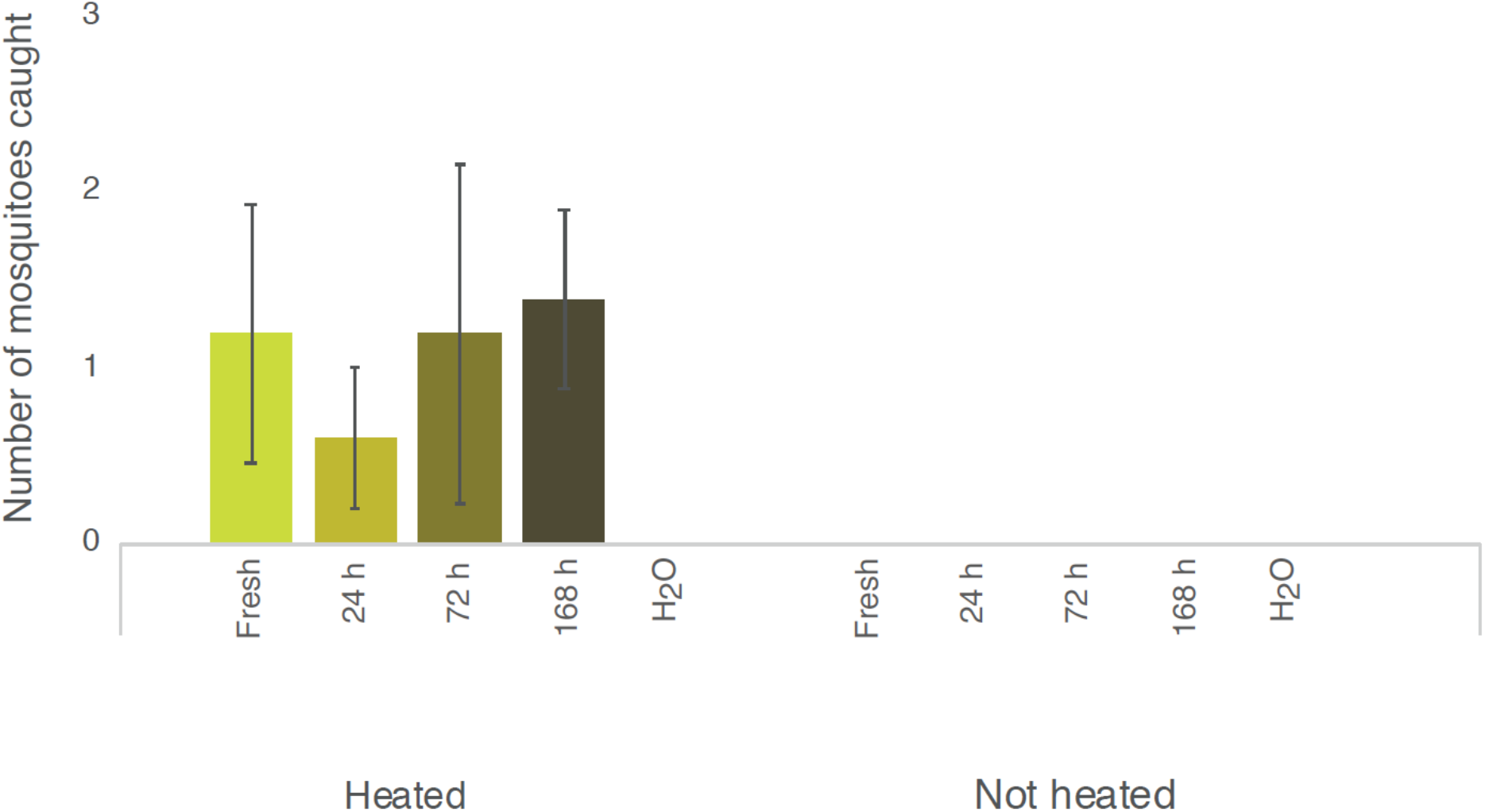
Cattle urine enhances host decoy trap catches only in the presence of the host cue, heat. Host decoy traps only caught malaria mosquitoes in a deserted pasture between the breeding site and the village in the presence of both heat and cattle urine (fresh or aged), but not either alone. Error bars indicate the standard error of the mean.

## Notes

### Competing Interest Statement

The authors have declared no competing interest.

